# A small DNA virus initiates replication with no more than three genome copies per cell

**DOI:** 10.1101/2022.05.27.493787

**Authors:** Ruifan Ren, Limin Zheng, Junping Han, Camila Perdoncini Carvalho, Shuhei Miyashita, Deyong Zhang, Feng Qu

**Affiliations:** Longping Branch Graduate School, Hunan University, Changsha, China; Department of Plant Pathology, The Ohio State University, Wooster, OH 44691, United States of America; Hunan Plant Protection Institute, Changsha, China; Graduate School of Agricultural Science, Tohoku University, Japan

**Keywords:** single-stranded DNA virus, intracellular virus population bottleneck, superinfection exclusion, virus evolution, natural selection

## Abstract

Cellular organisms purge lethal mutations as they occur (in haploids), or as soon as they become homozygous (in sexually reproducing diploids), thus making the mutation-carrying genomes the sole victims of lethality. How lethal mutations in viruses are purged remains an unresolved question because numerous viral genomes could potentially replicate in the same cell, sharing their encoded proteins, hence shielding lethal mutations from selection. Previous investigations by us and others suggest that viruses with plus strand (+) RNA genomes may compel such selection by bottlenecking the replicating genome copies in each cell to low single digits. However, it is unclear if similar bottlenecks also occur in cells invaded by DNA viruses. Here we investigated whether tomato yellow leaf curl virus (TYLCV), a small virus with a single-stranded DNA genome, underwent population bottlenecking in cells of its host plants. We engineered the TYLCV genome to produce two replicons that express green fluorescent protein and mCherry, respectively, in a replication-dependent manner. We found that less than 65% of cells penetrated by both replicons replicated both, whereas at least 35% of cells replicated either of them alone, illustrating an intracellular population bottleneck size of no more than three. Furthermore, sequential inoculations unveiled strong mutual exclusions of these two replicons in most cells. Collectively our data demonstrated for the first time that DNA viruses like TYLCV are subject to stringent intracellular population bottlenecks, suggesting that such population bottlenecks may be a virus-encoded, evolutionarily conserved trait that assures timely elimination of lethal mutations.

**Significance statement:** An important unresolved issue in virus life cycles is how natural selection acts on individual virus copies in the same cells. Unlike cellular organisms in which genome copies with lethal or advantageous mutations usually share their hosts with no more than one homologous genome copy, viruses could potentially reproduce with numerous sister genomes per cell, permitting sharing of protein products, thereby greatly diminishing phenotypic impacts of otherwise eventful mutations. Previous investigations suggest that (+) RNA viruses solve this problem by bottlenecking the number of replicating genome copies to low single digits. The current study reveals strikingly similar intracellular population bottlenecks for a DNA virus. Further mechanistic interrogations could avail the virus-encoded bottleneck-enforcing apparatus as targets for antiviral therapy and prevention.

## Introduction

The replication processes of many viruses are highly error prone, causing most descendant viral genomes to differ from their parents by a minimum of one mutation (1–5). While these mutations are mostly phenotypically neutral or near neutral, lethal mutations do occur and can reach high numbers, given that millions of new viral genomes are frequently produced in each cell. Viral genomes containing loss-of-function mutations within genes encoding proteins, especially proteins essential for viral replication, if not promptly isolated and purged, pose serious threat to the survival of the cognate virus population. This is because these “cheaters” could steal the corresponding functional proteins produced by sister genomes in the same cell to support their own reproduction. More ominously, absent of diligent surveillance, similar cheater mutants arise continuously as the replication reiterates in new cells. Together they could quickly reproduce themselves to dominance, and steadily dilute out the genomes still producing functional proteins, ultimately obliterating the cognate viral population (1, 2, 6, 7). Therefore, the law of natural selection predicts that successful viruses must have evolved ways to weed out lethal mutations in a timely manner. However, exactly how this is accomplished remains poorly understood.

Recent observations by us and others prompted a new hypothesis that explains how plus-strand (+) RNA viruses purge lethal mutations (8–11). Briefly, this Bottleneck, Isolate, Amplify, Select (BIAS) hypothesis postulates that multiple genome copies of the same virus, upon penetrating the same cell, cooperate to erect intracellular population bottlenecks from which very few copies (as few as one) of the viral genome could escape to initiate replication. Due to the stochastic nature of the bottlenecks, a viral genome copy with a lethal mutation has the same chance as other sister genomes to escape the bottlenecks and use replication proteins produced from sister genomes to initiate replication. However, should this occur, its replication would lead to the amplification of descendants that all contain the same lethal mutation. Upon invading a fresh cell collectively, such a mutant genome lineage would then be forced to bear the consequence of the lethal mutation, because the mutation-complementing sister genomes would now be absent (or inadequate if the bottleneck size is >1), leading to the elimination or drastic suppression of viral genome copies bearing lethal mutations.

While the BIAS hypothesis is amply supported by evidence derived from (+) RNA viruses, it is not known whether it also applies to DNA viruses. To address this knowledge gap, here we examined a small DNA virus to determine whether its populations also encountered narrow bottlenecks in infected cells. We adopted tomato yellow leaf curl virus (TYLCV), a worldwide pathogen of tomato, as the model virus for this study. TYLCV is a member of the genus *Begomovirus*, family *Geminiviridae*, with a single-stranded, circular DNA genome of approximately 2,800 nucleotides (nt), encoding six proteins (12–14). The four genes on the complementary strand of the genome are early expressing, encoding proteins (C1-C4) that participate in various aspects of viral genome replication, transcriptional activation, and host defense neutralization (Figure 1A) (13, 15). In particular, C1 is absolutely required for the rolling circle replication of TYLCV genome (16–18). On the other hand, the two genes on the viral strand of TYLCV genome are both late expressing, dependent on successful genome replication. They encode V1 and V2, which are capsid protein (CP) and suppressor of RNA silencing, respectively (Figure 1A). V1 (CP) is not essential for intracellular replication of TYLCV (19). In the current report, a TYLCV genome was modified to encode nuclearly localized fluorescent proteins in place of V1, permitting convenient tracking of viral replication. We found that TYLCV intracellular populations were constrained by stringent bottlenecks that limit replicating genome copies to no more than three in each cell. Furthermore, TYLCV genomes that entered cells early exerts strong superinfection exclusion (SIE) against those entering merely 24 hours later. Together these data suggest that DNA viruses like TYLCV may also utilize a strategy similar to BIAS to enable timely elimination of lethal mutations.

**Figure 1.**
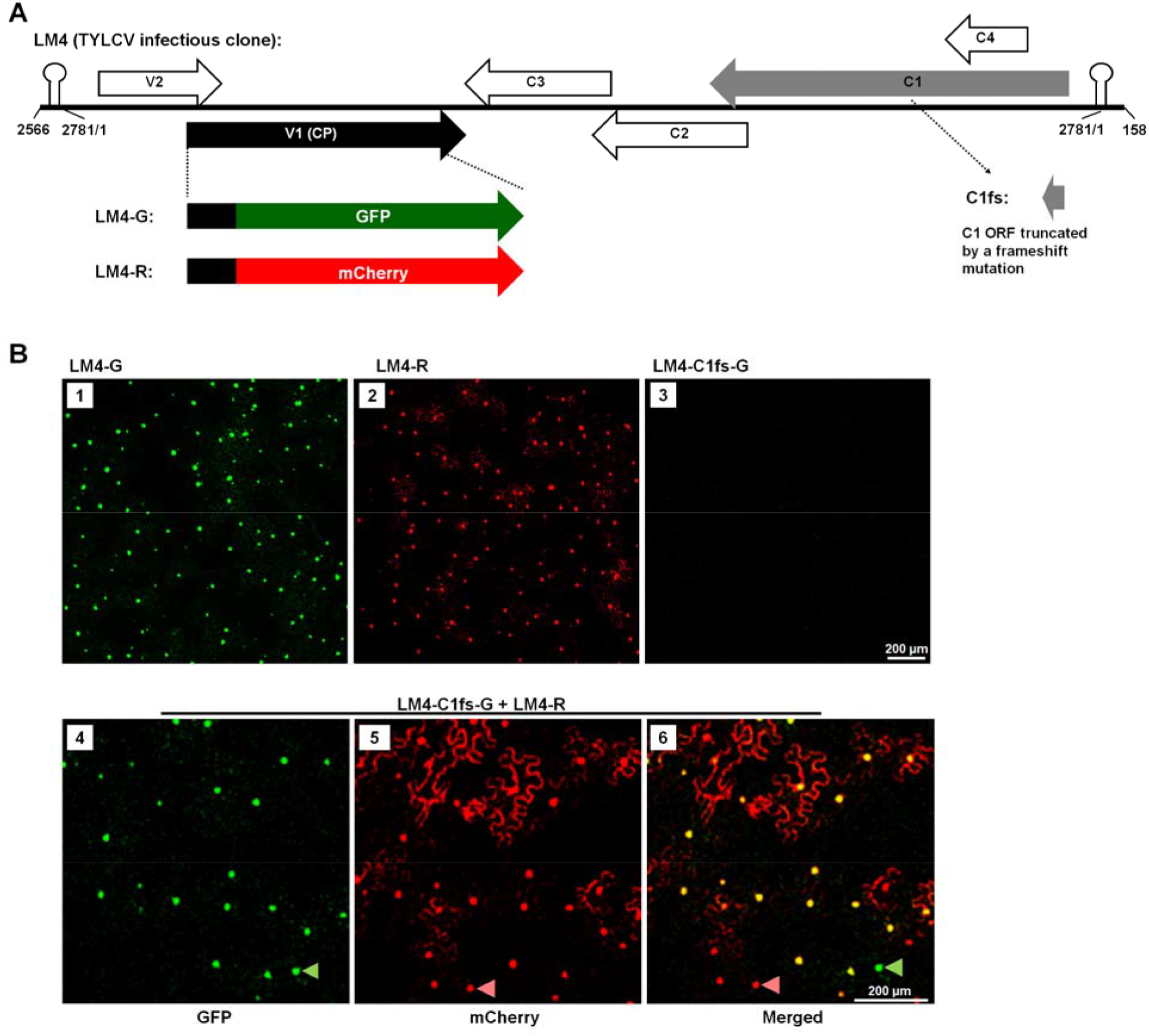
TYLCV-derived replicons LM4-G and LM4-R replicate to produce nuclearly localized fluorescent signals. **A**. Schematic depiction of the 2,781-nt TYLCV genome in linearized, double-stranded form inserted in a binary shuttle plasmid. Note that nt positions 2566-2781 (256 nt) and 1-158 (158 nt) were duplicated at 5’ and 3’ ends, respectively. The large arrows depict various TYLCV-encoded proteins. The V1 ORF, which were modified to accommodate GFP and mCherry insertions, is highlighted as solid black. The C1 ORF was also modified in one of the constructs (LM4-C1fs-G) and is highlighted as solid gray. **B**. Replication-dependent expression of GFP and mCherry from LM4-G and LM4-R, respectively. Bar = 200 μm.

## Results

### TYLCV as a model for examining intracellular population bottlenecks of DNA viruses

TYLCV is a small virus with a single-stranded, circular DNA genome of approximately 2,800 nt. Previous studies by others established that expression of the V1 gene of TYLCV, encoding viral CP, occurs in a replication-dependent manner (20–22). We hence reasoned that TYLCV replication could be monitored microscopically by replacing the CP coding sequence with that of green fluorescence protein (GFP) or the red fluorescent mCherry. Such replacements gave rise to two TYLCV replicons designated as LM4-G and LM4-R, respectively (Figure 1A). Note that the V1 and V2 genes partially overlap. As a result, the V2-overlapping portion of V1 was retained in LM4-G and LM4-R in the form of N-terminal fusions to GFP and mCherry. This 65 amino acid (aa) N-terminal fusion harbored a strong nuclear localization signal (NLS) that routed most of the GFP and mCherry signals to the cell nuclei, permitting convenient quantification of TYLCV-replicating cells.

To test their replicability, the LM4-G and LM4-R replicons were cloned into the pAI101 binary vector (23). The resulting constructs, still referred to as LM4-G and LM4-R for simplicity, were transformed into *Agrobacterium tumefaciens* (agro; strain C58C1), suspensions of which were pressure-infiltrated into leaves of *Nicotiana benthamiana* plants (agro-infiltration). Both replicons produced abundant, strongly fluorescent signals that were predominantly located in the nuclei of epidermal cells (Figure 1B, panels 1 and 2). As a control, we produced the LM4-C1fs-G mutant by deleting a single nt early in the C1 open reading frame (ORF) of LM4-G, causing the C1 frame to shift and stop prematurely. This mutant no longer produced any GFP fluorescence (Figure 1A; Figure 1B, panel 3), verifying that GFP expression from LM4-G indeed depended on viral replication. Importantly, the C1 defect in LM4-C1fs-G was successfully complemented by wildtype C1 proteins provided *in trans* from the co-delivered LM4-R replicon (Figure 1B, panels 4-6), demonstrating that these two replicons readily entered the same cells. Interestingly, cells that expressed just one of the two fluorescent proteins were also observed (Figure 1B, panels 4 – 6, green and purple arrowheads). While it might be expected that some cells replicated only LM4-R, it was surprising that cells exclusively replicating the defective replicon LM4-C1fs-G also existed (Figure 1B, panels 4-6, green arrowhead). Therefore, even though the replication-competent LM4-R provided C1 proteins to complement the replication of the defective replicon LM4-C1fs-G, its own replication in the same cell was not needed for complementation to occur. More informatively, this observation also revealed that not all C1-producing TYLCV genome copies were destined for replication in the cells they invaded, thus offering the earliest hint for the existence of population bottlenecks during TYLCV intracellular reproduction. Together the above experiments established LM4-G and LM4-R as robust TYLCV replicons suitable for examining the intracellular bottlenecking of TYLCV populations.

### Two TYLCV replicons mix-delivered through agro-infiltration enter same *N. benthamiana* cells at very high frequencies

To determine whether TYLCV populations became bottlenecked inside the cells they invaded, we first must ensure that cells infiltrated with the replicon-containing agro suspensions do internalize multiple copies of the TYLCV genome. Note that upon successful attachment, each agro cell was known to load each plant cell with at least six copies of the T-DNA (24). Accordingly, cells replicating LM4-G or LM4-R should already contain multiple copies of either replicon. Instead, what we want to ensure is the same-cell-internalization of two different, easily distinguishable replicons (LM4G and LM4-R).

Ideally detection of cells that replicate both LM4-G and LM4-R would indicate their same cell co-existence. However, existence of extremely narrow intracellular bottlenecks would limit the number of the internalized genome copies that could replicate to produce GFP and mCherry, making this indicator unreliable. Therefore, alternative approaches were needed to assure the same-cell entry of both replicons. The first approach we took was to count the number of fluorescent cells produced by LM4-G or LM4-R alone, each delivered at two different agro concentrations, with the respective OD_600_ values adjusted to 0.05 and 0.5. We reasoned that if these two agro doses, differing by 10 folds, produced similar numbers of fluorescent cells, then the lower OD 0.05 dose must be sufficient to infect most susceptible cells. Accordingly, the higher OD 0.5 dose would be predicted to load each of the cells with multiple copies of the replicon genome. To this end, six *N. benthamiana* leaves were infiltrated with LM4-G or LM4-R-containing agros, with the two halves of the same leaf receiving the same agro strain diluted to OD 0.05 and 0.5, respectively. For each replicon/concentration combination, we counted fluorescent cells in 10 viewing fields of 2.4 mm^2^ (1.55 mm X 1.55 mm) in size, randomly selected from six leaves. This experiment was repeated three times, with the results summarized in SI Dataset 1 and SI Figure 1A. While in the first two repeats the OD 0.5 concentration correlated with slightly more fluorescent cells, in the third repeat the LM4-R replicon at OD 0.5 actually produced slightly fewer fluorescent cells than at OD 0.05. However, the differences never exceeded 1.5 folds. Therefore, the lower OD 0.05 concentration must have been sufficient to deliver the replicons into at least 70% of susceptible cells. Accordingly, the OD 0.5 dilution was above saturation by at least five folds, and almost certain to guarantee the entry of both replicons into the same cells.

We further corroborated this conclusion by mixing replicon-containing agros (LM4-G, LM4-R) with those containing constructs designed to express GFP or mCherry without viral replication (35S-GFP and 35S-mCherry). Note that for this set of experiments all agro suspensions were diluted to OD 0.1 before mixing, meaning the final concentration for each agro strain was OD 0.05. Furthermore, in the control experiment where 35S-GFP and 35S-mCherry were mixed, another agro strain harboring a construct encoding the p19 protein of tomato bushy stunt virus was needed to counteract RNA silencing (25), hence the final concentration for each of the agro strains was OD 0.033. SI Figure 1B showed that at this concentration, the replication-independent expression of GFP and mCherry occurred in almost all cells, thus both fluorescence co-existed in nearly 100% of the cells. Similarly, when the non-replicating 35S-GFP was mixed with the replicating LM4-R, or 35S-mCherry with LM4-G, the agro concentration of OD 0.05 each was sufficient to cause nearly all cells to produce both GFP and mCherry (SI Figure 1C and 1D). Together these data confirmed that agros at the concentration of OD 0.05 were enough to (i) cause nearly all treated cells to express GFP or mCherry independent of virus replication; and (ii) permit two or more co-delivered constructs to enter the same cells at near 100% frequency. Thus, mixed delivery of LM4-G and LM4-R via agro-infiltration would be expected to achieve similar levels of same-cell penetration even with the agro concentration of OD 0.05.

### LM4-G and LM4-R co-replicate in less than 65% of cells that receive both constructs

We next used the LM4-G and LM4-R replicons to assess whether multiple TYLCV genome copies entering the same cells could all initiate replication. Mixture of agro suspensions harboring the two replicons was again diluted to two different concentrations (OD 0.05 and 0.5 for each of the replicons), and applied to two sides of the same leaf. The infiltrations were done on at least six leaves. At four days after infiltration, 10 2.4-mm^2^ viewing fields were randomly chosen for quantification of cells that replicated LM4-G, LM4-R, or both. Furthermore, the entire procedure was repeated three times to ensure reproducibility. Figure 2A shows typical confocal images of one viewing field collected under different light channels. Comparison of images of different channels (GFP, mCherry, Merged) permitted straightforward differentiation of cell nuclei that expressed GFP or mCherry, or both. Figure 2B shows the quantification results of more than 10,000 cells. Since the total numbers of cells with fluorescent nuclei varied from leaf to leaf, we converted the counts of each viewing field to percentages to permit side-by-side comparisons. As shown in Figure 2B and SI Dataset 2, across the six repeats (three at OD 0.05 and three at OD 0.5), the percentage means for cells replicating solely LM4-G ranged from 15.7% to 20.1%, whereas those replicating only LM4-R ranged from 22.6 to 30.1%. Finally, cells replicating both replicons were between 49.8 and 60.9%. These data unveiled a consistent trend of stringent intracellular population bottlenecking that limit the number of TYLCV genome copies replicating in each cell to low single digits (more in next section). Note that in mixed infections, LM4-G consistently replicated in a slightly smaller number of cells that LM4-R (SI Dataset 2; the relative LM4-G/LM4-R ratio (g : r) was calculated for each repeat/concentration combination and listed on the top of the tables). This small difference between cells replicating LM4-G and LM4-R translated into a mean ratio of 47 : 53, which were factored into subsequent estimation of bottleneck sizes.

**Figure 2.**
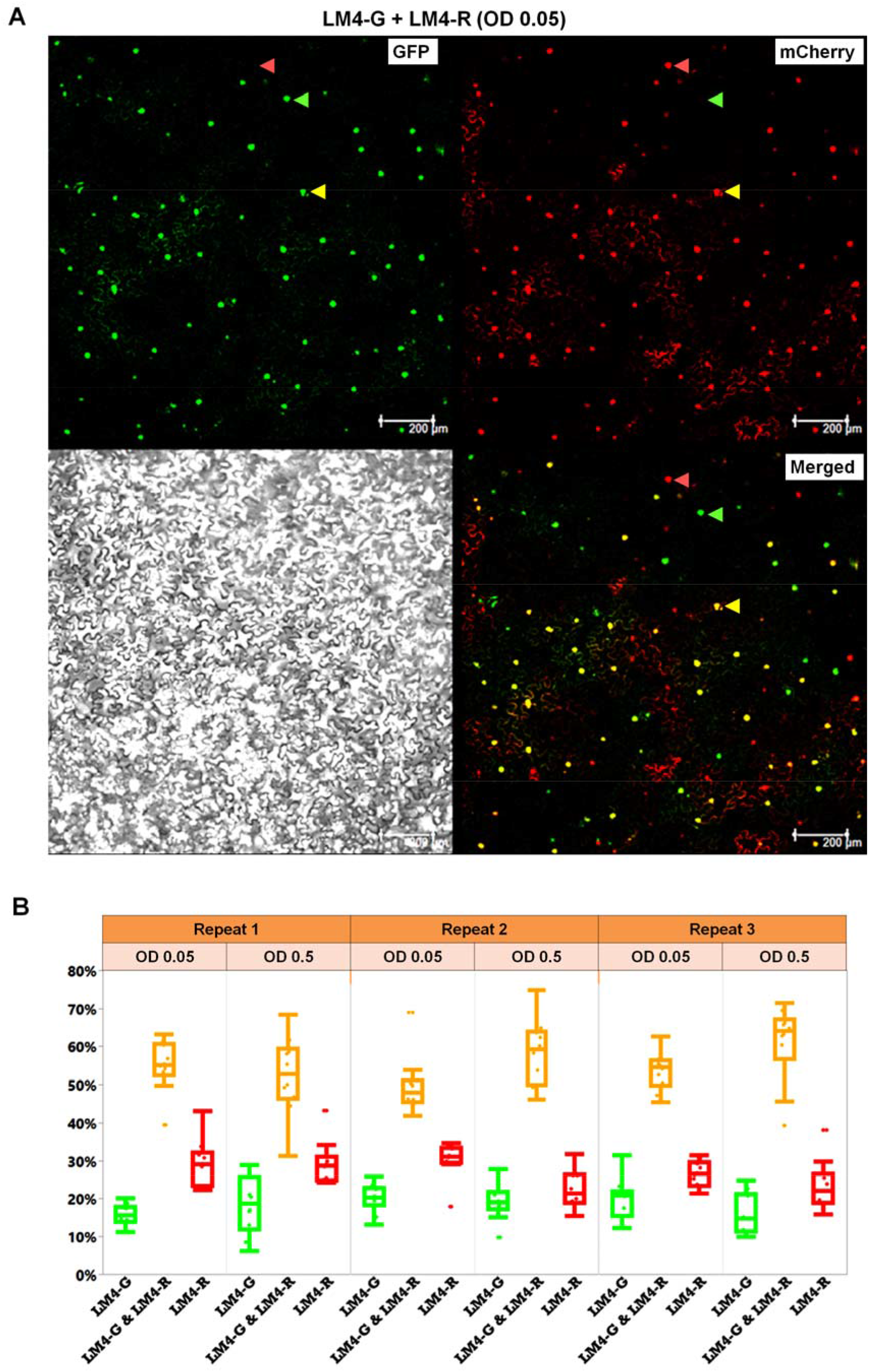
Revelation of TYLCV intracellular population bottlenecks with the LM4-G and LM4-R replicons. **A**. Images of a typical 2.4 mm^2^ viewing field taken from a leaf section that received both LM4-G and LM4-R, showing nuclei that emit green (GFP) or red (mCherry) fluorescence, or both (Merged), one of each highlighted with arrowheads of different colors. The gray scale image serves as reference for cell sizes, shapes, and boundaries. **B**. Quantification of percentages of cells replicating LM4-G, LM4-R, or both (LM4-G & LM4-R). Shown are box plots derived from numeration data of six experimental groups (three repeats, each with two different agro concentrations, 10 data points per group).

### The size of TYLCV intracellular population bottlenecks is no larger than three

To gauge the size of TYLCV intracellular population bottlenecks, we considered the initiation of replication by two or more replication-competent replicons in the same cell as independent events. This is then analogous to blindly sampling a jar filled with green (g) and red (r) jellybeans, one at a time. Assuming the jar contains an equal number of green and red beans, the chance of getting green or red beans with one sampling attempt is (g + r)/2, equaling to 50%, whereas the chance of getting both beans is zero. Repeating the sampling one more time, the chances of getting green, red, or both beans can be expressed as (g + r)/2 X (g + r)/2, equaling to (g^2^ + 2gr + r^2^)/4, meaning there is a 25% chance of getting either green or red beans (g^2^ or r^2^), and a 50% chance (2gr) of getting both beans. The chance predictions for sampling up to four times are summarized in SI Table 1. As the sampling time increases to more than four times, the chance of getting both beans increases to at least 87.5%, whereas the chance of getting solely green or red beans dwindles to below 6.25%.

We now get back to the data with LM4-G and LM4-R replicons, borrowing the symbols g and r to represent LM4-G and LM4-R, respectively. The observed percentage means (SI Dataset 2; 15.7 – 20.1% for GFP only, 49.8 – 60.9% for both GFP and mCherry, and 22.6 – 30.1% for mCherry only) fit nicely with the predicted outcomes of two sampling attempts, suggesting that the bottleneck size was around 2. To obtain more precise estimates, we assumed that the variation of bottleneck sizes among individual cells obeyed Poisson distribution, with an average size of *λ*. We then computed *λ* and its standard deviation using a maximum-likelihood method (see Materials and Methods for details). Additionally, since replication of LM4-G alone occurred in slightly fewer cells than LM4-R alone, we simultaneously estimated the ratio of g relative to (g + r) in each repeat to account for the effect of replication bias. We obtained *λ* values of 2.38 ± 0.05 and 2.60 ± 0.05 for OD 0.05 and 0.5, respectively (SI Table 2), with the g/(g + r) ratio ranging from 0.42 to 0.48. This suggests that the size of intracellular TYLCV population bottlenecks was no more than three, and was largely unaffected by a 10-fold increase of inoculum dose.

### TYLCV intracellular population bottlenecks remain stringent when the LM4-G and LM4-R replicons are delivered with one combined construct

To further rule out the possibility that GFP-only or mCherry-only cells resulted from failure of both replicons to invade the same cells, we next assembled a new construct, designated LM4-G_LM4-R, by combining both replicons in the same DNA molecule. The LM4-G_LM4-R construct was then delivered into *N. benthamiana* cells at two agro concentrations (OD 0.05 and 0.5), and cell nuclei expressing GFP only, both, or mCherry only were counted for three repeat experiments. As shown in Figure 3 and SI Dataset 3, this new construct appeared to further compromise the relative competitiveness of LM4-G. This notion was supported by maximum-likelihood estimations, which showed that the g/(g + r) ratio was significantly lower than 0.5 (varying from 0.13 ± 0.01 to 0.32 ± 0.01; SI Table 3). Nevertheless, the estimated bottleneck sizes, 2.37 ± 0.07 and 2.20 ± 0.08 for OD 0.05 and 0.5, respectively, were not substantially different from the estimates obtained with two separate constructs. Thus, guaranteeing the same cell co-existence of both LM4-G and LM4-R by combining them in the same construct provided further corroboration of earlier estimation of an intracellular bottleneck size of no more than three.

**Figure 3.**
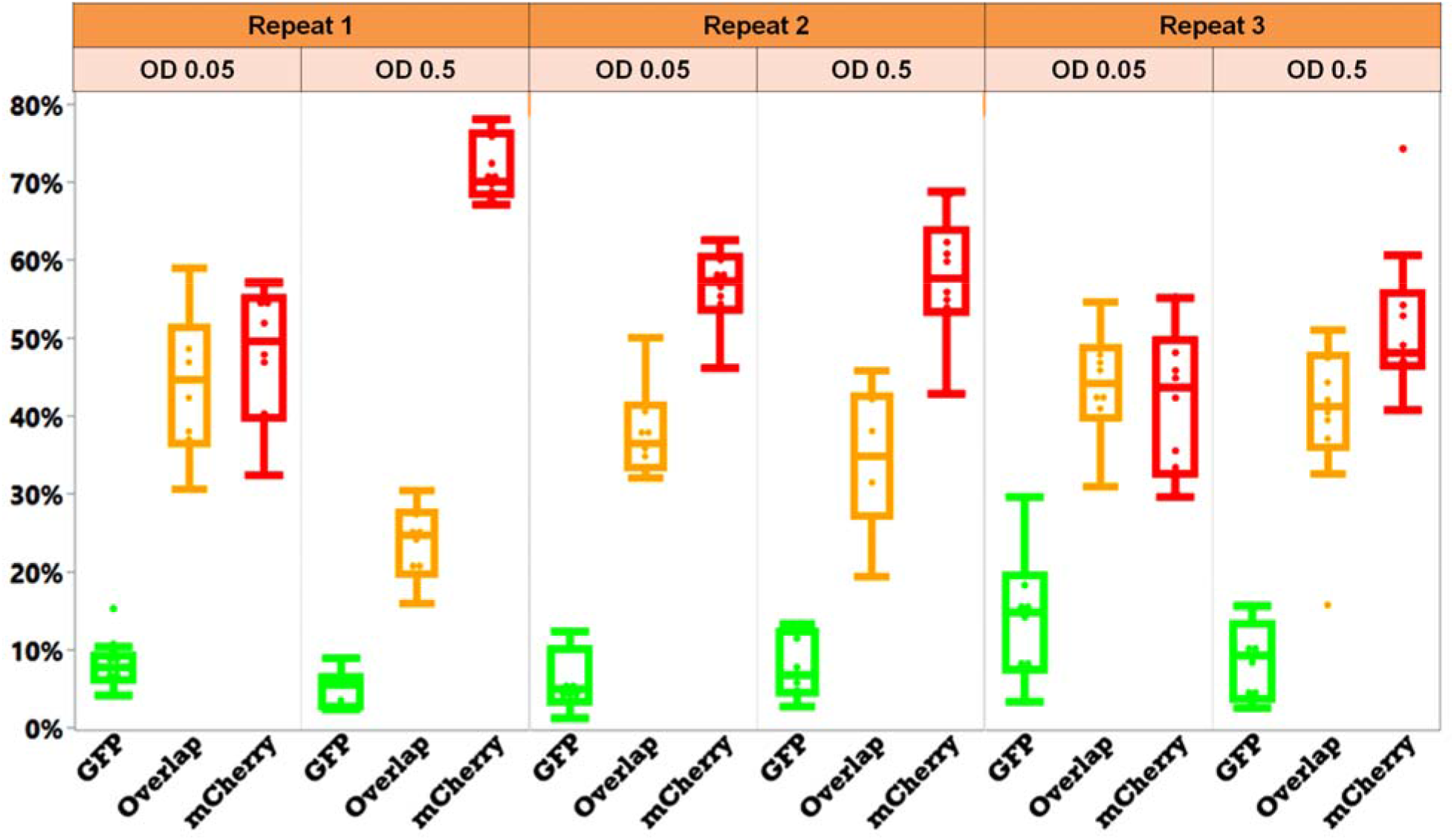
Quantification of percentages of cell nuclei replicating LM4-G, LM4-R, or both (LM4-G & LM4-R) in leaves treated with one construct that carry both replicons (LM4G_LM4R). Shown are box plots derived from numeration data of six experimental groups (three repeats, each with two different agro concentrations, 10 data points per group).

### LM4-G and LM4-R exert strong intracellular SIE to each other

We reported earlier that in (+) RNA virus infections SIE manifested intracellular bottlenecking of viral populations (8–11, 26, 27). To determine whether TYLCV also exhibited intracellular SIE, we next used the LM4-G and LM4-R replicons to infect *N. benthamiana* leaf cells in a sequential manner. As controls, *N. benthamiana* leaves pre-infiltrated with either infiltration buffer or the replication-independent 35S-GFP construct was unable to block the replication of the superinfecting LM4-R (Figure 4 A and B). Furthermore, prior entry of the replication-defective LM4-C1fs-G construct not only failed to block LM4-R replication, but actually permitted the latter to complement the defect, leading to co-replication of both in many, but not all cells (Figure 4C). By contrast, prior invasion of the replication-competent LM4-G replicon conditioned a strong SIE against LM4-R, blocking the latter from replicating in most cells (Figure 4D). Indeed, among more than 1,000 cells inspected in multiple repeat experiments, nearly all replicated only the pre-delivered LM4-G, with only one cell found to replicate both (Figure 4D, yellow arrowhead). Conversely, prior invasion of LM4-R completely blocked the replication of the superinfecting LM4-G (Figure 4E and F). Therefore, LM4-G and LM4-R mutually excluded each other in cells in which their own replication was established.

**Figure 4.**
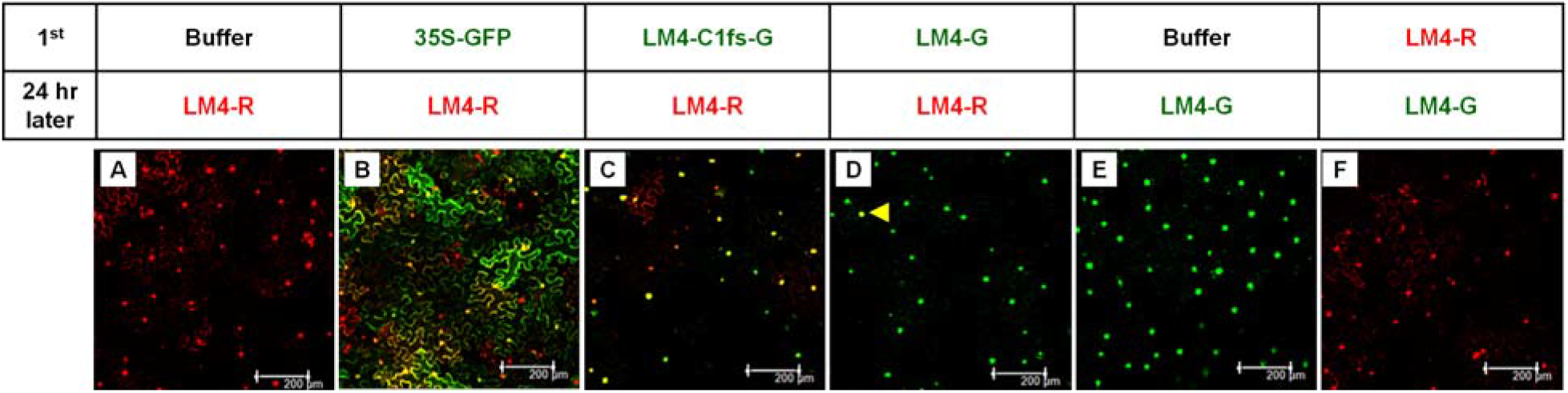
LM4-G and LM4-R exhibits mutual exclusion at the cellular level when delivered sequentially with a 24 hour (hr) interval.

## Discussion

In the current study, we sought to resolve whether copies of a DNA virus genome entering the same cell could all initiate replication or, whether they are subject to intracellular reproductive bottlenecking in a manner similar to (+) RNA viruses (9, 10). To this end, two TYLCV replicons, LM4-G and LM4-R, were created that expressed two different fluorescent proteins (GFP and mCherry) in a replication-dependent manner. These two replicons were then introduced into leaves of *N. benthamiana* plants in the form of pre-mixed agro suspensions to ensure both entered the same cells. Additionally, they were also delivered in the form of a single plasmid construct (LM4-G_LM4-R) to guarantee their simultaneous penetration of the same cells. Importantly, GFP and mCherry expressed from LM4-G and LM4-R replicons predominantly translocated to the cell nuclei, permitting straightforward numeration of cells that replicated one or both replicons. Counting of tens of thousands of fluorescent cell nuclei, followed by probability computation, led to the conclusion that TYLCV populations were severely bottlenecked intracellularly, permitting no more than three genome copies to commence replication in each cell.

Exactly how many genome copies of a virus initiate replication in a single cell is a critical question because it is becoming increasingly clear that most viruses invade most cells with not one, but many virions, hence many copies of viral genomes (28). Consistent with this view, copies of the (+) sense poliovirus genomic RNA were found to direct translation of viral proteins at multiple intracellular sites before commencing replication (29). On the other hand, even if just one copy of viral genome entered a cell initially and it succeeded in launching replication, we are still faced with the question of how many progeny genome copies repeat the replication cycle in the cell of their parent.

The need to address this question is even more evident considering the high error rate of the viral replication processes, estimated to be approximately 10^−4^ for every nt incorporated (7, 30, 43). Depending on the size of viral genomes, this error rate translates into almost one mutation for every new genome copy synthesized. Given the fact that viruses often replicate millions of progeny genome copies in each infected cell, it is inevitable that some of the progeny genome copies contain loss-of-function mutations in essential protein-coding genes. If even 100 of the progeny genomes were permitted to re-replicate in the cell of their own genesis, those with lethal errors would be able to hitchhike on the functional proteins produced by sister genomes, thus could not be efficiently purged from the virus population.

While so far we have emphasized the critical importance of weeding out lethal mutations to the survival of virus populations, also worth noting is that absent of bottleneck-mediated isolation, a viral genome copy with an advantageous mutation in a protein-coding gene would not be the only one, or even the primary one, to benefit from the encoded advantage. Absent of this positive feedback, advantageous mutations would have no way to become more numerous in the virus population. To put it differently, positive selection could not occur. Conversely, the abundance of literature documenting potent purifying as well as positive selections acting on virus populations suggests that most viruses must have evolved ways to subject mutations to timely selection (32–35). The BIAS arrangement would be one of the ways to meet this end, thus itself can be expected to be positively selected.

Besides serving as a DNA virus model for the BIAS hypothesis, the TYLCV system has additional advantages. To address the challenge of constant emergence of viral mutations, earlier researchers proposed that some of the replication proteins encoded by (+) RNA viruses serve exclusively the very RNA from which they are translated, hence are cis-acting (36, 37). The cis-acting hypothesis poses serious evolutionary problems, but is difficult to rule out completely for (+) RNA viruses for which both translation and replication occur in the cytoplasm. By contrast, the cis-acting arrangement would be impossible to achieve for DNA viruses like TYLCV. This is because the TYLCV-encoded replication protein (C1) must be translated in the cytoplasm of the infected cells, and then routed back to cell nuclei to orchestrate the rolling circle replication. It is then impossible to differentiate between C1 proteins originated from different TYLCV genome copies.

In summary, with the current study we have demonstrated for the first time that populations of small DNA viruses like TYLCV are intracellularly bottlenecked in a manner similar to (+) RNA viruses. This finding provides exciting leads for follow-up studies aimed at elucidating the mechanism(s) of the bottlenecking, and assessing its potential role in facilitating natural selection in viruses. Outcomes of these follow-up studies will likely avail the bottlenecking machinery as the target for preventive as well as therapeutic interventions of virus diseases of plants, animals, and humans.

## Materials and Methods

### Constructs

The original TYLCV infectious clone (Y10. The Genbank accession number of the corresponding TYLCV SH2 isolate is AM282874.1) was kindly provided by Dr. Xueping Zhou of China Institute of Plant Protection (38). The full-length, double-stranded form of TYLCV genome, plus a 216-bp duplication at the 5’ end, and a 158-bp duplication at the 3’ end, was subcloned into pAI101, a *E.coli-A. tumefaciens* shuttle vector modified from pCambia1300 in our lab (23, 39), leading to a new TYLCV infectious clone we call LM4. To create LM4-G and LM4-R, two KpnI sites were introduced into the V1 gene, at positions 307/308 and 881/882 (numbering relative to the full length genome sequence), respectively. The coding sequences of uvGFP and mCherry were then PCR amplified and cloned between the KpnI site using the NEBuilder kit (New England Biolabs). The sequences of LM4, LM4-G and LM4-R encompassing the entire TYLCV DNA (and its modified forms) were verified with Sanger sequencing. Other constructs (p19, 35-GFP, 35S-mCherry) were described previously (27, 40, 41).

### Agrobacterium infiltration (agro-infiltration)

All DNA constructs destined for testing in *N. benthamiana* plants were transformed into electrocompetent A. tumefaciens strain C58C1 via electroporation using the AGR setting on the Bio-Rad Micropulser Electroporator. Briefly, 5 μl of the plasmid DNA was mixed with 40 μl of agro cells and maintained on ice until electroporation. After electroporation, 900 μl of SOB media was added and the suspension was incubated at 28 °C for one hour. Selection was carried out on solid Terrific Broth (TB) media containing rifampicin, gentamycin, and kanamycin. Successful introduction of the plasmid was confirmed using colony PCR. A single colony confirmed to have the desired plasmid was used to inoculate 3 ml TB liquid media with the same antibiotics, and incubated overnight at 28 °C. The culture was diluted 1:100 with fresh TB liquid media and incubated under the same conditions for another night. The second culture was centrifuged at 4,000 rpm for 20 min, and resuspended in agroinfiltration buffer (10 mM MgCl2, 10 mM MES, and 100 μM acetosyringone). All suspensions were diluted to OD600 =1 and incubated at 28 °C for 3 hours. *Agrobacterium* suspensions were then mixed and introduced into leaves of young *N. bethamiana* plants via a small wound, using a needleless syringe.

### Confocal microscopy

Four days after agro-infiltration, leaf discs were collected from the plants. Confocal microscopy was performed at the Molecular and Cellular Imaging Center (MCIC), the Ohio Agricultural Research and Development Center, using a Leica DMI6000 laser confocal scanning microscope. To detect GFP and mCherry fluorescence, sequential excitation at 488 nm and 587 nm was provided by argon and helium-neon 543 lasers, respectively.

### Numeration of the fluorescent nuclei

To count the cells that replicate LM4-G, LM4-R, or both, images of 2.4 mm^2^ (1.55 mm X 1.55 mm) were collected using a 10X lens, from randomly selected leaf sections receiving different combinations of agro suspensions. For samples treated with the mixture of LM4-G and LM4-R, or the combined LM4G_LM4-R construct, images of three separate channels were collected for every selected viewing field to allow for the separate counting of different colored spots (GFP, mCherry, or all spots). For each treatment group, at least 10 images were collected from six different leaves. The fluorescent spots representing nuclei of infected cells were counted using the ImageJ program. The number of nuclei that simultaneously replicated both LM4-G and LM4-R was calculated by subtracting the number of all spots from the sum of green and red spots.

### Statistics and bottleneck size computation

The calculation of sums, means, percentages, standard deviations, were mostly carried out with various tools available through Excel. The box plots were generated with the JMP Pro 16.0.0 software package. Bottleneck size estimations were performed using a maximum likelihood algorithm described in our previous studies (8, 9, 41), based on the R software package ver. 4.1.3 (42). The script is available as Supplementary Text 1, and downloadable at GitHub (https://github.com/ShuheiMiyashita/Ren_et_al_2022).

## Acknowledgements

We thank Dr. Xueping Zhou of China Institute for Plant Protection for generously sharing the Y10 infectious clone. We are indebted to Dr. K. Andrew White for critically reading the manuscript, and making insightful suggestions. Members of the Qu lab are greatly appreciated for discussions and technical assistances. This study is supported in part by the NSF grant 1758912. RFR is supported in part by a scholarship from China Scholarship Council, and the Hunan Graduate Scientific Research Innovation Project (QL20210118). LMZ is supported in part by a grant from the National Key R&D Program of China (2018YFE0112600). SM is supported in part by Japan Society for the Promotion of Science (JSPS) KAKENHI grant 21K05591.

## Supplementary Information (SI)

The following SI files are available online:

SI dataset 1 – 3 as Excel files.

A combined SI appendix that includes:

- SI Text, an R script for *λ* and g/(g + r) computation – pages 2 and 3;
- SI Figure 1 with legends – page 4;
- SI Tables 1 – 3, page 5.

## References

1. M. Eigen, Error catastrophe and antiviral strategy. Proc. Natl. Acad. Sci. U.S.A. 99, 13374– 13376 (2002).

2. J. J. Bull, R. Sanjuán, C. O. Wilke, Theory of lethal mutagenesis for viruses. J. Virol. 81, 2930–2939 (2007).

3. R. Sanjuán, P. Agudelo-Romero, S. F. Elena, Upper-limit mutation rate estimation for a plant RNA virus. Biol. Lett. 5, 394–396 (2009).

4. J. M. Malpica, et al., The rate and character of spontaneous mutation in an RNA virus. Genetics 162, 1505 (2002).

5. R. French, D. C. Stenger, Evolution of wheat streak mosaic virus: Dynamics of population growth within plants may explain limited variation. Annu. Rev. Phytopathol. 41, 199–214 (2003).

6. E. Domingo, C. Perales, Viral quasispecies. PLoS Genet. 15, e1008271 (2019).

7. S. Duffy, Why are RNA virus mutation rates so damn high? PLoS Biol. 16, e3000003– e3000003 (2018).

8. S. Miyashita, H. Kishino, Estimation of the size of genetic bottlenecks in cell-to-cell movement of soil-borne wheat mosaic virus and the possible role of the bottlenecks in speeding up selection of variations in trans-acting genes or elements. J. Virol. 84, 1828– 1837 (2010).

9. S. Miyashita, K. Ishibashi, H. Kishino, M. Ishikawa, Viruses roll the dice: The stochastic behavior of viral genome molecules accelerates viral adaptation at the cell and tissue levels. PLoS Biol. 13, e1002094 (2015).

10. X.-F. Zhang, et al., A self-perpetuating repressive state of a viral replication protein blocks superinfection by the same virus. PLoS Path. 13, e1006253 (2017).

11. F. Qu, et al., Bottleneck, Isolate, Amplify, Select (BIAS) as a mechanistic framework for intracellular population dynamics of positive-sense RNA viruses. Virus Evol. 6 (2020).

12. A. Prasad, N. Sharma, G. Hari-Gowthem, M. Muthamilarasan, M. Prasad, Tomato yellow leaf curl virus: Impact, challenges, and management. Trends Plant Sci. 25, 897–911 (2020).

13. L. Hanley-Bowdoin, E. R. Bejarano, D. Robertson, S. Mansoor, Geminiviruses: masters at redirecting and reprogramming plant processes. Nat. Rev. Microbiol. 11, 777–788 (2013).

14. X. Zhou, Advances in understanding begomovirus satellites. Annu. Rev. Phytopathol. 51, 357–381 (2013).

15. V. N. Fondong, Geminivirus protein structure and function: Geminivirus proteins. Mol. Plant Pathol. 14, 635–649 (2013).

16. B. M. Orozco, L. Hanley-Bowdoin, Conserved sequence and structural motifs contribute to the DNA binding and cleavage activities of a geminivirus replication protein. J. Biol. Chem. 273, 24448–24456 (1998).

17. B. M. Orozco, A. B. Miller, S. B. Settlage, L. Hanley-Bowdoin, Functional domains of a geminivirus replication protein. J. Biol. Chem. 272, 9840–9846 (1997).

18. B. M. Orozco, L.-J. Kong, L. A. Batts, S. Elledge, L. Hanley-Bowdoin, The multifunctional character of a geminivirus replication protein Is reflected by its complex oligomerization properties. J. Biol. Chem. 275, 6114–6122 (2000).

19. Hanley-Bowdoin L, Elmer J S, Rogers S G, Expression of functional replication protein from tomato golden mosaic virus in transgenic tobacco plants. Proc. Natl. Acad. Sci. 87, 1446– 1450 (1990).

20. G. Sunter, D. M. Bisaro, Transactivation of geminivirus AR1 and BR1 gene expression by the viral AL2 gene product occurs at the level of transcription. Plant Cell 4, 1321 (1992).

21. G. Sunter, D. M. Bisaro, Transactivation in a geminivirus: AL2 gene product is needed for coat protein expression. Virology 180, 416–419 (1991).

22. G. Sunter, D. M. Bisaro, Regulation of a geminivirus coat protein promoter by AL2 protein (TrAP): Evidence for activation and derepression mechanisms. Virology 232, 269–280 (1997).

23. J. Lin, et al., The Bean pod mottle virus RNA2-encoded 58-kilodalton protein P58 is required in cis for RNA2 accumulation. J. Virol. 88, 3213–3222 (2014).

24. H. Oltmanns, et al., Generation of backbone-free, low transgene copy plants by launching T-DNA from the Agrobacterium chromosome. Plant Physiol. 152, 1158–1166 (2010).

25. F. Qu, T. J. Morris, Efficient infection of Nicotiana benthamiana by Tomato bushy stunt virus Is facilitated by the coat protein and maintained by p19 through suppression of gene silencing. Mol. Plant-Microbe Interact. 15, 193–202 (2002).

26. X.-F. Zhang, et al., Random plant viral variants attain temporal advantages during systemic infections and in turn resist other variants of the same virus. Sci. Rep. 5 (2015).

27. Q. Guo, et al., Superinfection exclusion by p28 of turnip crinkle virus Is separable from its replication function. Mol. Plant-Microbe Interact. 33, 364–375 (2020).

28. R. Sanjuán, Collective infectious units in viruses. Trends Microbiol. 25, 402–412 (2017).

29. D. Egger, K. Bienz, Intracellular location and translocation of silent and active poliovirus replication complexes. J. Gen. Virol. 86, 707–718 (2005).

30. S. F. Elena, R. Sanjuán, Adaptive value of high mutation rates of RNA viruses: separating causes from consequences. J. Virol. 79, 11555–11558 (2005).

31. J. T. McCrone, et al., Stochastic processes constrain the within and between host evolution of influenza virus. eLife 7, e35962 (2018).

32. M. D. Pauly, M. C. Procario, A. S. Lauring, A novel twelve class fluctuation test reveals higher than expected mutation rates for influenza A viruses. eLife 6, e26437 (2017).

33. N. D. Grubaugh, et al., Genetic drift during systemic arbovirus infection of mosquito vectors leads to decreased relative fitness during host switching. Cell Host & Microbe 19, 481–492 (2016).

34. N. D. Grubaugh, Translating virus evolution into epidemiology. Cell Host & Microbe 30, 444–448 (2022).

35. K. Kawamura-Nagaya, K. Ishibashi, Y.-P. Huang, S. Miyashita, M. Ishikawa, Replication protein of tobacco mosaic virus cotranslationally binds the 5’ untranslated region of genomic RNA to enable viral replication. Proc. Natl. Acad. Sci. 111, E1620–E1628 (2014).

36. G. Yi, K. Gopinath, C. C. Kao, Selective repression of translation by the brome mosaic virus 1a RNA replication protein. J. Virol. 81, 1601 (2007).

37. H. Zhang, H. Gong, X. Zhou, Molecular characterization and pathogenicity of tomato yellow leaf curl virus in China. Virus Genes 39, 249–255 (2009).

38. S. Zhang, R. Sun, Q. Guo, X.-F. Zhang, F. Qu, Repression of turnip crinkle virus replication by its replication protein p88. Virology 526, 165–172 (2019).

39. F. Qu, T. Ren, T. J. Morris, The coat protein of turnip crinkle virus suppresses posttranscriptional gene silencing at an early initiation step. J. Virol. 77, 511–522 (2003).

40. R. Sun, S. Zhang, L. Zheng, F. Qu, Translation-independent roles of RNA secondary structures within the replication protein coding region of turnip crinkle virus. Viruses 12, 350 (2020).

41. D. A. Abebe, S. van Bentum, M. Suzuki, S. Ando, H. Takahashi, S. Miyashita, Plant death caused by inefficient induction of antiviral R-gene-mediated resistance may function as a suicidal population resistance mechanism. Communications Biology 4, 947 (2021).

42. R Core Team, R: A language and environment for statistical computing. R Foundation for Statistical Computing, Vienna, Austria. URL https://www.R-project.org/ (2022).

43. S. Duffy, E.C. Holmes, Phylogenetic evidence for rapid rates of molecular evolution in the single-stranded DNA begomovirus tomato yellow leaf curl virus. J. Virol. 82, 957–965 (2008).

