## Supplementary material for "A small DNA virus initiates replication with no more than three genome copies per cell": Combined Supplementary Information (from another version of this preprint: version 1, 2022-05-27)

**The combined SI appendix includes:**

- SI Text, an R script for $\lambda$ and g/(g + r) computation – pages 2 and 3;
- SI Figure 1 with legends – page 4
- SI Tables 1 – 3, page 5.

**SI Text – an R script for $\lambda$ and g/(g + r) computation**

```
#An R script for bottleneck size estimation

### Data input
# cgo: no. of cells with GFP only
# cgr: no. of cells with both GFP and mCherry
# cro: no. of cells with mCherry only
# data for separate-vector system (OD 0.05) are used as an example
cgo1 <- 200; cgr1 <- 701; cro1 <- 373 # Rep1
cgo2 <- 336; cgr2 <- 834; cro2 <- 498 # Rep2
cgo3 <- 175; cgr3 <- 468; cro3 <- 231 # Rep3


### Main body of bottleneck size estimation

K <- 30 #maximum number of founders included in the calculation
kv <- NULL
lv <- NULL
for (i in 0:K){
kv <- c(kv,rep(i,i+1))
lv <- c(lv,0:i)
}
klv <- kv-lv
lklv <- lv*klv
ln <- (K+2)*(K+1)/2

## Function for log likelihood for any lambda (bottleneck size) and r (ratio of g to g+r)
FmLL <- function(lr){
lambda <- lr[1]; r1 <- lr[2]; r2 <- lr[3]; r3 <- lr[4];
t <- matrix(rep(0,ln*7),ncol=7)
t[,1] <- kv
t[,2] <- lv
t[,3] <- klv
t[,4] <- lklv
t[,5] <- dpois(t[,1],lambda)*dbinom(t[,2],t[,1],r1)
t[,6] <- dpois(t[,1],lambda)*dbinom(t[,2],t[,1],r2)
t[,7] <- dpois(t[,1],lambda)*dbinom(t[,2],t[,1],r3)

#Rep1
pni1 <- t[1,5]
pgo1 <- sum(t[which(t[,2]>0&t[,3]==0),5])/(1-pni1)
pro1 <- sum(t[which(t[,2]==0&t[,3]>0),5])/(1-pni1)
pgr1 <- 1-pgo1-pro1

#Rep2
pni2 <- t[1,6]
pgo2 <- sum(t[which(t[,2]>0&t[,3]==0),6])/(1-pni2)
```

```
pro2 <- sum(t[which(t[,2]==0&t[,3]>0),6])/(1-pni2)
pgr2 <- 1-pgo2-pro2

#Rep3
pni3 <- t[1,7]
pgo3 <- sum(t[which(t[,2]>0&t[,3]==0),7])/(1-pni3)
pro3 <- sum(t[which(t[,2]==0&t[,3]>0),7])/(1-pni3)
pgr3 <- 1-pgo3-pro3

#Log likelihood
LL1 <- dmultinom(c(cgo1,cgr1,cro1),prob=c(pgo1,pgr1,pro1),log=TRUE)
LL2 <- dmultinom(c(cgo2,cgr2,cro2),prob=c(pgo2,pgr2,pro2),log=TRUE)
LL3 <- dmultinom(c(cgo3,cgr3,cro3),prob=c(pgo3,pgr3,pro3),log=TRUE)
LL <- LL1 + LL2 + LL3

-LL
}

## Maximization of log likelihood to find most likely lambda and r
init <- c(5,0.5,0.5,0.5)
FmLL.opt <- optim(init,FmLL, NULL, method="L-BFGS-B", hessian = TRUE, lower=c(0.1,0.01,0.01,0.01),
upper=c(10,0.99,0.99,0.99))
FmLL.opt$par  # lambda and r estimates
v <- solve(FmLL.opt$hessian)
sd <- sqrt(diag(v))
sd        # standard deviations for lambda and r, respectively
```

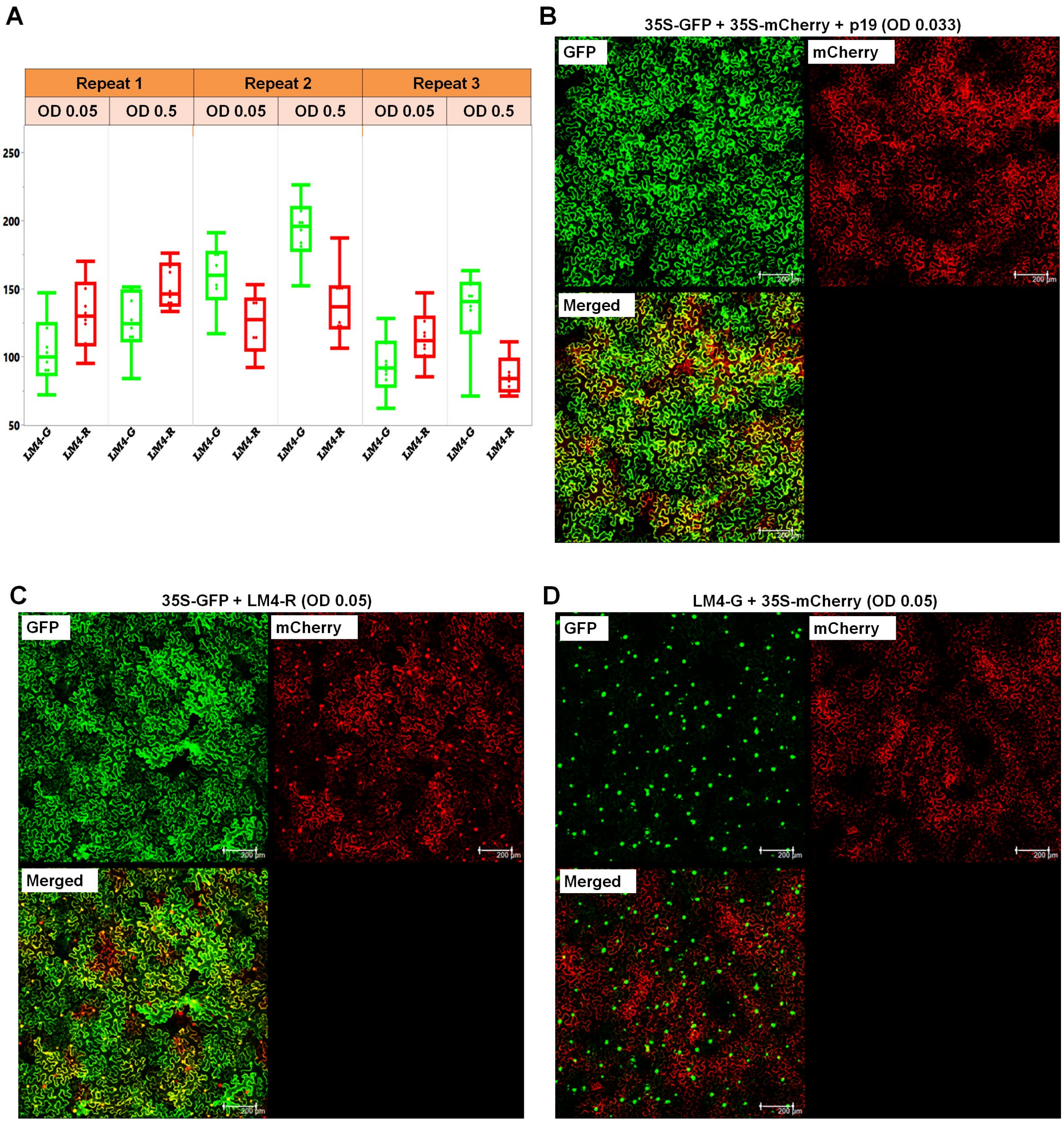


**SI Figure 1**. Entry of multiple TYLCV replicons into the same *N. benthamiana* cells via agroinfiltration delivery. **A**. The number of cells replicating LM4-G or LM4-R when they were delivered separately, at two different agro concentrations (OD 0.05 and OD 0.5). Note the numbers on Y-axle were actual counts of fluorescent cell nuclei. Each of the box plots reflect 10 randomly selected viewing fields of 2.4 $mm^2$. **B**. Cells agro-infiltrated with the non-replicating 35-GFP and 35S-mCherry, at OD 0.033, serving as control for **C** and **D**. **C**. Near complete overlap of the fluorescence signals produced by the non-replicating 35S-GFP and the replicating LM4-R, both delivered with agro concentration at 0.05. **D**. Similar to **C**, shown here is the near complete overlap of the fluorescence signals produced by the replicating LM4-G and the non-replicating 35S-mCherry, both delivered with agro concentration at 0.05.

**SI Table 1. Formulas for estimating bottleneck sizes**

| Bottleneck size | Formula (assuming g and r are present at 1:1 ratio) | Predicted outcomes | | |
|---|---|---|---|---|
| | | green | both | red |
| 1 | $[(g + r)/2]^1 = 0.5g + 0.5r$ | 50% | 0 | 50% |
| 2 | $[(g + r)/2]^2 = 0.25g^2 + 0.5gr + 0.25r^2$ | 25% | 50% | 25% |
| 3 | $[(g + r)/2]^3 = 0.125g^3 + 0.375g^2r + 0.375gr^2 + 0.125r^3$ | 12.5% | 75% | 12.5% |
| 4 | $[(g + r)/2]^4$<br>$= 0.0625g^4 + 0.25g^3r + 0.375g^2r^2 + 0.25gr^3 + 0.0625r^4$ | 6.25% | 87.5% | 6.25% |

**SI Table 2. Maximum-likelihood estimates of *λ* and g/(g + r) with LM4-G and LM4-R as separate constructs**

| | ***λ*** | **g/(g + r)** | | |
|---|---|---|---|---|
| | | Repeat 1 | Repeat 2 | Repeat 3 |
| Estimates (OD=0.05) | 2.38 ± 0.05 | 0.42 ± 0.01 | 0.44 ± 0.01 | 0.46 ± 0.01 |
| Estimates (OD=0.5) | 2.60 ± 0.05 | 0.43 ± 0.01 | 0.48 ± 0.01 | 0.45 ± 0.01 |

**SI Table 3. Maximum-likelihood estimates of *λ* and g/(g + r) with LM4-G and LM4-R combined in a single construct**

| | ***λ*** | **g/(g + r)** | | |
|---|---|---|---|---|
| | | Repeat 1 | Repeat 2 | Repeat 3 |
| Estimates (OD=0.05) | 2.37 ± 0.07 | 0.27 ± 0.01 | 0.21 ± 0.01 | 0.32 ± 0.01 |
| Estimates (OD=0.5) | 2.20 ± 0.08 | 0.13 ± 0.01 | 0.22 ± 0.01 | 0.26 ± 0.01 |
